# Phosphorylation of β-arrestins impacts recruitment to Parathyroid Hormone Receptor 1 (PTH1R) at the cell surface and endosomes

**DOI:** 10.64898/2026.10.05.756893

**Authors:** Emma R Carruthers, Caitlin R M Oyagawa, Raewyn C Poulsen, Natasha L Grimsey

## Abstract

β-arrestins 1 and 2 are intracellular adaptor proteins that regulate the subcellular localisation and signalling of GPCRs including Parathyroid Hormone Receptor 1 (PTH1R), which produces β-arrestin-dependent signalling that may selectively stimulate bone anabolism. β-arrestins can themselves be regulated by post-translational modifications including phosphorylation, which dictates the translocation and signalling of the few GPCRs studied in this context to date. We aimed to investigate whether phosphorylation of β-arrestins modulates their subcellular translocation in response to PTH1R activation. Translocation of β-arrestins to the plasma membrane and early endosomes was quantified using BRET biosensors and immunocytochemistry with high content analysis. Phosphoknockout or phosphomimetic mutations were introduced to previously demonstrated phosphorylation sites in β-arrestins 1 and 2, initially in bulk to assess overall effects. Plasma membrane translocation was altered only subtly, whereas early endosome translocation was markedly reduced for both mutations but to a greater degree for the phosphomimetic. Contributions of individual or paired sites were assessed by restoration to wildtype from the bulk phosphomimetic mutants. Dual restoration of S13 and T275 in β-arrestin 1, or the analogous sites in β-arrestin 2, restored the greatest degrees of early endosomal translocation, with complete restoration for β-arrestin 1 and partial for β-arrestin 2. Restoration of other sites had minimal or no effect. Dephosphorylation therefore appears to be required for PTH1R-stimulated β-arrestin early endosomal translocation, with different phosphorylation sites implicated than some other GPCRs. These findings raise the possibility of differential β-arrestin regulation by distinct stimuli which may have implications in therapeutic targeting of β-arrestin-dependent pathways.

**Graphical Abstract:** 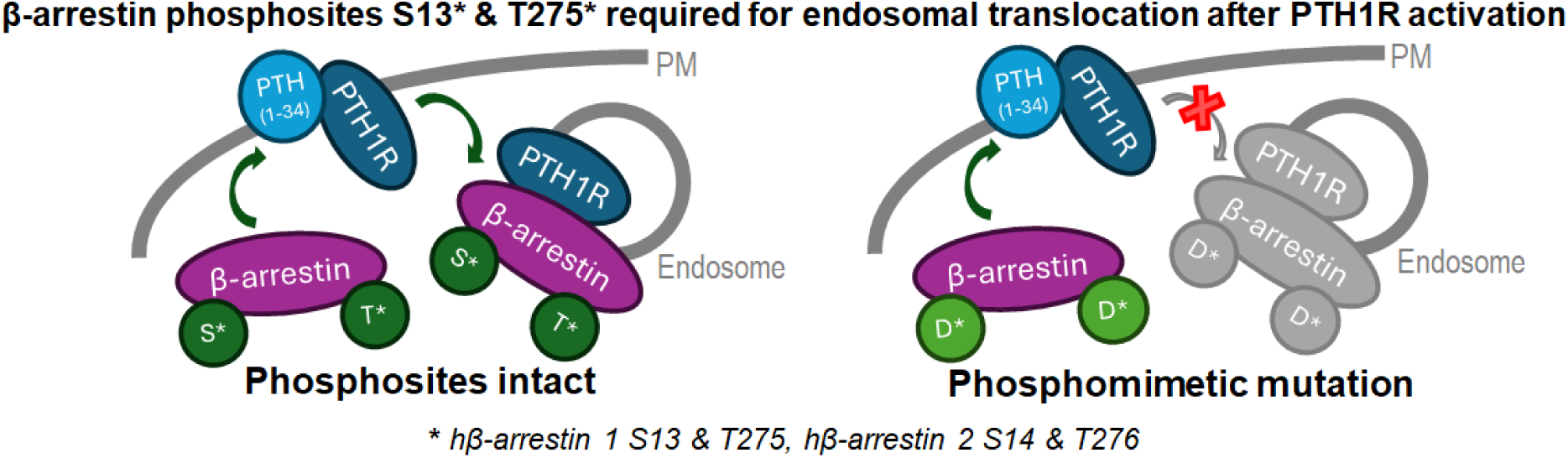

## Introduction

β-arrestins 1 and 2 (also known as arrestins 2 and 3) are intracellular adaptor proteins that act as key regulators of G protein-coupled receptor (GPCR) subcellular localisation and signalling. Mechanisms of this regulation include attenuation of acute GPCR signalling by desensitisation and promotion of receptor internalisation (Smith & Rajagopal, 2016), and formation of scaffolds with signalling proteins to activate pathways such as MAPK (Peterson & Luttrell, 2017). The critical importance of GPCR cytoplasmic domain phosphorylation (or presence of phospho-mimetic residues) in GPCR-arrestin interactions is extremely well established, with the barcode hypothesis describing how the pattern of GPCR phosphorylation directs the conformation in which β-arrestins bind and their subsequent downstream functions (Mayer et al., 2019). While less widely studied to date, the distribution and function of β-arrestins can also be influenced by a range of post-translational modifications on β-arrestins themselves, including phosphorylation (Table 1).

**Table 1.** Empirically demonstrated phosphorylation sites in β-arrestins 1 and 2, mutated in pKO and pMim in this study. ^a^ Residue numbering corresponds to human arrestin sequences

| β-arrestin | Site<sup>a</sup> | GPCR studied | Phosphorylation effect on GPCR internalisation | Kinase | Basal state | Reference(s) |
| --- | --- | --- | --- | --- | --- | --- |
| 1 | S13 | NA (predicted site) |  | ERK |  | Paradis et al. (2015) (predicted) |
|  | T275 | NA (predicted site) |  | ERK |  | Paradis et al. (2015) (predicted) |
|  | T374 | 5-HT4 |  |  |  | Cassier et al. (2017) |
| | S412 | β2AR<sup>1-3</sup>, 5-HT4<sup>4</sup> | Inhibits | ERK<sup>2</sup>, GRK5<sup>4</sup> | Phosphorylated | <sup>1</sup> Lin et al. (1997) ; <sup>2</sup> Lin et al. (1999); <sup>3</sup> Luttrell et al. (1999); <sup>4</sup> Barthet et al. (2009) |
| 2 | S14 | CXCR4, V2R, AT1R | Promotes | ERK |  | Paradis et al. (2015) |
|  | S194 | 5-HT2C, 5-HT4 |  |  |  | Cassier et al. (2017) |
|  | S264 | CXCR4, V2R, AT1R | No effect | ERK |  | Paradis et al. (2015) |
|  | S266/<br>S267 | 5-HT2C, 5-HT4 |  |  |  | Cassier et al. (2017) |
|  | T276 | CXCR4, V2R, AT1R | Promotes | ERK |  | Paradis et al. (2015) |
|  | S280 | 5-HT2C, 5-HT4 |  |  |  | Cassier et al. (2017) |
| | S360 | β2AR<sup>5</sup>, 5-HT2C<sup>6</sup>, 5-HT4<sup>6</sup> | Inhibits | | Phosphorylated | <sup>5</sup> Lin et al. (2002) ; <sup>6</sup> Cassier et al. (2017) |
| | T382 | β2AR<sup>5</sup>, 5-HT2C<sup>6</sup> | Inhibits<sup>5</sup>, no effect<sup>7,6</sup> | Casein kinase 2<sup>5,7</sup>, MEK<sup>6</sup> | Phosphorylated | <sup>5</sup> Lin et al. (2002); <sup>6</sup> Cassier et al. (2017); <sup>7</sup> Kim et al. (2002) |

For β-arrestin 1, a well-established site is serine (S) 412 (numbering for human β-arrestin 1), which is basally phosphorylated and becomes dephosphorylated upon β2 adrenergic receptor (β2AR) activation, facilitating receptor internalisation and binding to various signalling proteins (Lin et al., 1997; Luttrell et al., 1999). The main sites identified in β-arrestin 2 are likewise basally phosphorylated, with dephosphorylation upon activation resulting in receptor internalisation. These include S14 and T276 (Paradis et al., 2015), which may exert synergistic effects on β-arrestin 2 conformation (Kang et al., 2020). Analogous sites are present in β-arrestin 1 (S13 and T275), mutation of which induced similar conformational changes as the corresponding sites in β-arrestin 2 (Kang et al., 2020), but to date have not been demonstrated experimentally to undergo phosphorylation. Potentially conflicting findings exist for β-arrestin 2 T382, with one study finding agonist-induced dephosphorylation of T382 and S360 promoted β2AR internalisation (Lin et al., 2002), but another finding no effect on internalisation (Kim et al., 2002). Upon activation of a different GPCR, 5-HT_2c_, T383 (the rat orthologue equivalent of human T382) exhibited increased phosphorylation resulting in ERK activation but no effect on receptor internalisation (Cassier et al., 2017). This finding specifically highlights the potential for differential phosphorylation patterns with different stimuli.

Parathyroid Hormone Receptor 1 (PTH1R) is a GPCR involved in bone metabolism and serum mineral ion homeostasis. Stimulation with parathyroid hormone (PTH) elicits rapid and sustained recruitment of β-arrestins 1 and 2 to PTH1R at the plasma membrane, followed by co-internalisation of β-arrestins and receptor into endosomes (Ferrari et al., 1999). These endosomal PTH1R-β-arrestin complexes can generate prolonged cAMP (Ferrandon et al., 2009) and pERK (Gesty-Palmer et al., 2006; Sneddon et al., 2007) signalling. There is evidence that β-arrestin-biased PTH1R signalling via these pathways can selectively stimulate anabolic pathways in bone while avoiding the concurrent catabolic effects exerted by balanced ligands such as PTH, which may be therapeutically beneficial in osteoporosis (Gesty-Palmer et al., 2011). To date, effects of β-arrestin phosphorylation in response to PTH1R activation have not been studied.

Given the phosphorylation state of specific sites in β-arrestins 1 and 2 can regulate their trafficking and function in response to activation of non-PTH1R GPCRs, we hypothesised that phosphorylation downstream of PTH1R activation could also affect β-arrestin function. This study aimed to investigate the influence of phosphoknockout and phosphomimetic mutation on β-arrestin translocation in response to PTH1R activation. Translocation to the cell surface and early endosomes were quantified using a bystander NanoBRET assay employing subcellularly-targeted BRET acceptors, supplemented with immunocytochemistry. Previously reported phosphorylation sites were mutated in combination and individually to narrow down roles of specific sites.

## Methods

### β*-arrestin Expression Plasmids and Cloning*

Human β-arrestins 1 and 2 tagged at the N-terminus with Nanoluciferase (Nluc) in pcDNA3 were kindly provided by Michelle Glass and Jamie Manning, University of Otago (Manning et al., 2023).

Multi-phosphomutant β-arrestin coding sequences were purchased as pre-synthesised GenTitan^TM^ Gene Fragments (GenScript). Phosphoknockout (pKO) sequences were produced by exchanging the indicated (Table 1) wildtype human β-arrestin serine (S) or threonine (T) codons for alanine (A; GCC), whereas phosphomimetics (pMim) replaced native putative phosphosites with aspartic acid (D; GAC). Mutated β-arrestin sequences were cloned into pcDNA3-Nluc.

Vector backbones for assembly were amplified by PCR using KAPA HiFi HotStart polymerase (Roche). After PCR, methylated template DNA was removed by incubation with DpnI (New England Biolabs) for 60 minutes at 37°C. Products were gel-extracted using the Zymoclean Gel DNA Recovery Kit (Zymo). Assembly between PCR-amplified vector backbone and commercially synthesised gene fragments was performed using the NEBuilder HiFi DNA Assembly kit (New England Biolabs), in 2 μL reactions including 20 ng of the largest fragment and a 2x molar ratio of the smaller fragment. Products were transformed into XL 10 Gold Ultracompetent Cells (Agilent) and clonal colonies isolated and expanded. DNA was purified using the QIAprep Spin Miniprep Kit (Qiagen). Final constructs were verified by Sanger sequencing (Auckland Genomics, University of Auckland).

Single and double site ‘restoration’ mutants were generated by site-directed mutagenesis to re-introduce the wildtype serine or threonine codon at the indicated positions, using Nluc-β-arrestin 1 and 2 pMim multi-phosphomutant constructs as templates. Primers were designed according to Quikchange (Agilent) guidelines and synthesised (Integrated DNA Technologies). Primers were combined with β-arrestin pMim multi-phosphomutant plasmid template, KAPA HiFi HotStart polymerase, and appropriate buffer with nucleotides and cycled using the recommended conditions. Template was digested using DpnI as for seamless cloning. Products were transformed into XL 10 Gold Ultracompetent Cells and purified using the QIAprep Spin Miniprep Kit. Final constructs were verified by Sanger sequencing (Auckland Genomics, University of Auckland). Double site ‘restoration’ mutants were generated through a second round of mutagenesis using the relevant single ‘restoration’ mutants as templates.

### BRET β-arrestin Translocation

BRET β-arrestin translocation was performed essentially as previously described (Sharma et al., 2023), with minor modifications. HEK-293S cells (provided by Professor David Poyner, Aston Research Centre; Wootten et al., 2013) were maintained in DMEM (Cytiva, SH30243) with 8% foetal bovine serum (Moregate, New Zealand origin) at 37°C in a humidified atmosphere containing 5% CO_2_. Cells were plated at 40,000 cells per well in Poly-D-Lysine-treated 96-well white plates (Corning).

Cells were transfected with 45 ng/well hPTH1R (untagged in pcDNA3.1+; cDNA Resource Center, PTHR100000), 1 ng/well Nluc-β-arrestin, and 40 ng/well Venus-KRAS (plasma membrane marker; last 25 amino acids of human KRAS tagged with monomeric Venus) or Venus-Rab5 (early endosomal marker; human Rab5a tagged with monomeric Venus) using PEIMax® (Polysciences). Venus subcellular marker plasmids were kindly provided by Kevin Pfleger, University of Western Australia (Tiulpakov et al., 2016). PEIMax® (stock 0.05 mg/mL in PBS, 0.36 µL per well) and DNA were diluted separately in serum-free DMEM, then combined in equal volumes. Mixes were incubated for 60 minutes at room temperature before application of 40 µL per well to already-plated cells. Positions of transfections and stimulations (below) within 96-well plates were randomised between independent experiments.

Approximately 48 hours after transfection, cells were washed once with Hanks’ Balanced Salt Solution with calcium and magnesium (HBSS; Thermo Fisher Scientific 14025) + 1 mg/mL bovine serum albumin (BSA; low endotoxin and low free fatty acid; ICP Bio) (assay medium), then equilibrated in fresh assay medium for 30 minutes. Coelenterazine h (NanoLight Technology) was applied to cells in the dark from an intermediate dilution to reach a final concentration of 5 μM. Plates were read on a LUMIstar Omega (BMG Labtech) at 37°C using 475nm (Nluc) and 535nm (Venus) filters, initially for 5 minutes to monitor baseline. hPTH(1-34) (teriparatide acetate; Chemenu) was then applied from intermediate dilutions prepared at 5x the final intended concentration, and plates were read for 60 minutes.

BRET ratios were calculated as Venus / Nluc. Mean BRET ratios were calculated over 0-5 or 0-15 minutes as indicated and normalised as described in figure captions. Data with clear abnormalities, such as low raw Nluc counts relative to other replicates, were excluded from analysis. In the restoration phosphomutant experiment set, mean raw Nluc and Venus over the time period of interest from untransfected cells (as an indicator of background) were subtracted from the raw values of interest prior to calculating the BRET ratios.

### Immunocytochemistry (ICC) and High Content Image Analysis

All BRET experiments in this study had ICC plates prepared in parallel. Cells were plated and transfected as for BRET, using the same transfection mixes. hPTH(1-34) stimulations were carried out as close to simultaneously as possible, using the same aliquots of diluted ligand. Parallel ICC plates were stimulated for 15 minutes before fixation. At the conclusion of ligand stimulation, cells were fixed in paraformaldehyde (PFA, Electron Microscopy Sciences; 4% in phosphate buffer) for 10 minutes, followed by 2 washes in PBS-T (phosphate buffered saline with 0.2% Triton X-100 [Sigma-Aldrich]).

All antibodies were centrifuged at 14,000 xg for 10 minutes prior to use. Primary antibody anti-β-arrestin (BD Biosciences, 610551) was diluted 1:400 in immunobuffer (IB; PBS-T + 0.4 mg/mL merthiolate [BDH] and 1% goat serum [Sigma]), and incubated with rocking either for 3 hours at room temperature, or overnight at 4°C. Following incubation, cells were washed once with PBS-T. Secondary goat anti-mouse Alexa-647 (Thermo Fisher Scientific) was prepared as for the primary antibody and incubated on cells under the same conditions.

Nuclei were labelled with Hoescht 33258 in PBS-T (Sigma) for 15-30 minutes. Two final washes in PBS-T were performed then cells were stored in PBS-T with 0.4 mg/mL merthiolate.

Images were acquired on an ImageXpress Micro XLS Widefield High-Content Analysis System (Molecular Devices), across 9 sites per well using a 20x objective lens. Typically ∼1500-1800 HEK cells were captured in each image. Filters were DAPI for Hoechst, and Cy5 for Alexa 647. After analysis of individual images (see below), means for all sites in a well were taken, then the mean from technical replicate wells was calculated to produce the means ± SD for a single independent experiment. Occasional individual images with profound debris or abnormality relative to other images in the same well (e.g. out of focus, very low cell count) were excluded prior to data collation.

Quantification was performed using MetaXpress (Molecular Devices, v6.5.5.559) with automated analysis to detect membrane-like staining (continuous linear/tubular structures, expected to represent cell surface-localised β-arrestin) and punctate staining (expected to represent clustered and/or endosomal β-arrestin), building on prior published analyses (Finlay et al., 2016; Grimsey et al., 2008). In initial image processing, any bright debris was excluded by applying an intensity-based threshold mask on the staining-of-interest image, with dilation to ensure out-of-focus blur from the debris was also removed. The same “debris mask” area was removed from the corresponding nuclei staining image. Cell nuclei were counted on the Hoechst staining image using the “FindSpots” module. To identify the area of the image with detectable arrestin staining (including diffuse, membrane-like, and punctate), an intensity threshold was applied to the arrestin image to segment staining from background (as judged by comparison to negative controls, e.g. untransfected cells, and/or boundary between distinct patterns correlating to cell bodies versus diffuse signals between cells) and the pixel area within the threshold recorded (converted to μm^2^ via the image calibration). Prior to detection of membrane-like or punctate features, a blurred image (fast Fourier transform with “Blur” filter) at 50% intensity was subtracted from the original image, which had the effect of bringing the disparate staining intensities between cells into closer alignment within each image, thereby improving the veracity of subsequent feature detection. “Unsharp Mask” and “Median” filters were applied, which assisted in enhancing contrast for membrane-like staining. The “Angiogenesis” module (designed to detect areas of tube-like continuous staining) with “Skeletonization” algorithm was then applied to determine the image area corresponding to membrane-like staining. Prior to proceeding to puncta detection, the membrane-like staining area was dilated slightly and this area subtracted from the image to avoid any potential for bright membrane-like staining being re-detected as puncta. Punctate staining was quantified using the “Transfluor” module, which detects granular and/or spot-like image features as smaller puncta (“Pits”, 1-2 μm width) and larger puncta (“Vesicles”, 2-3 μm width). Data were analysed as membrane or total (small plus large) puncta area per cell, normalised to baseline raw Nluc signal (measured during BRET detection), then normalised to a control condition for collation of independent experiment data. The ratio of the area of small puncta to area of large puncta within each image was also calculated.

### Data and Statistical Analysis

Graphs were generated and statistical analysis was conducted using GraphPad Prism (v10, GraphPad Software). Data were normalised as described in figure captions, usually to the relevant time-matched vehicle condition (100%). As specified in figure captions, data are presented as the mean ± SD of two technical replicates or the mean ± SD of three or more independent experiments.

When statistical testing was of interest, one-or two-way analyses of variance (ANOVA) with repeated measures design were conducted on raw data (even where data is presented as normalised). Residual, homoscedasticity and QQ plots were visually inspected to determine fulfilment of parametric test assumptions of normality and equality of variance. If these plots indicated violation of assumptions, data were transformed (log10) and plots were re-inspected. Following transformation, parametric test assumptions were found to be met. If ANOVA indicated significant differences (p<0.05), Holm-Šídák post-hoc tests were performed on comparisons of interest.

## Results

### Mutation of β-arrestin phosphosites impacts recruitment to plasma membrane and endosomes as measured by BRET following PTH1R stimulation

To obtain overall indications of whether phosphorylation or dephosphorylation of β-arrestin affects its subcellular translocation upon stimulation of hPTH1R with hPTH(1-34), selected phosphorylation sites (Table 1) were simultaneously mutated to either non-phosphorylatable alanine (phosphoknockout, pKO) or aspartic acid which mimics phosphorylated serine or threonine (phosphomimetic, pMim). This mutation approach is established in studies of protein phosphorylation (Chen & Cole, 2015) including of β-arrestins (e.g. Paradis et al., 2015). Sites of interest were selected based on previous functional evidence of involvement downstream of other GPCR activation (Table 1). Wildtype and phosphomutant β-arrestins tagged with Nluc were expressed in HEK-S cells also expressing hPTH1R. BRET acceptors were Venus targeted to the cell surface (Venus-KRAS) or early endosomes (Venus-Rab5) to enable monitoring of β-arrestin translocation to both locations without requiring tagging of the receptor, an approach analogous to that adopted in previous studies (e.g. Tiulpakov et al., 2016).

When assessing arrestin proximity to the cell surface-targeted BRET acceptor Venus-KRAS, no difference in basal BRET was observed between wildtype and phosphomutant β-arrestins (Figure 1 A, B). Upon stimulation with 1 μM hPTH(1-34), both wildtype β-arrestins 1 and 2 demonstrated rapidly increased BRET ratios (indicating increased proximity to the plasma membrane), reaching maximum BRET ratios within ∼7 min. The β-arrestin 1 BRET signal then remained relatively stable to 60 minutes, with only a small steady decrease back toward baseline over the time course. This time-dependent decrease was observed to a slightly larger degree for β-arrestin 2. For both β-arrestins 1 (Figure 1A) and 2 (Figure 1B), pKO and pMim mutants displayed generally comparable plasma membrane translocation extents and temporal profiles to wildtype. Responses of β-arrestin 1 phosphomutants in particular were very similar to wildtype. Greater divergence was observed with β-arrestin-2, where pKO had a comparatively higher BRET ratio after the initial response, whereas pMim reached a similar plateau to wildtype but with slower onset during the first 5 minutes. These trends were also reflected in the corresponding β-arrestin 1 pKO and pMim, although to a much smaller degree.

**Figure 1.**
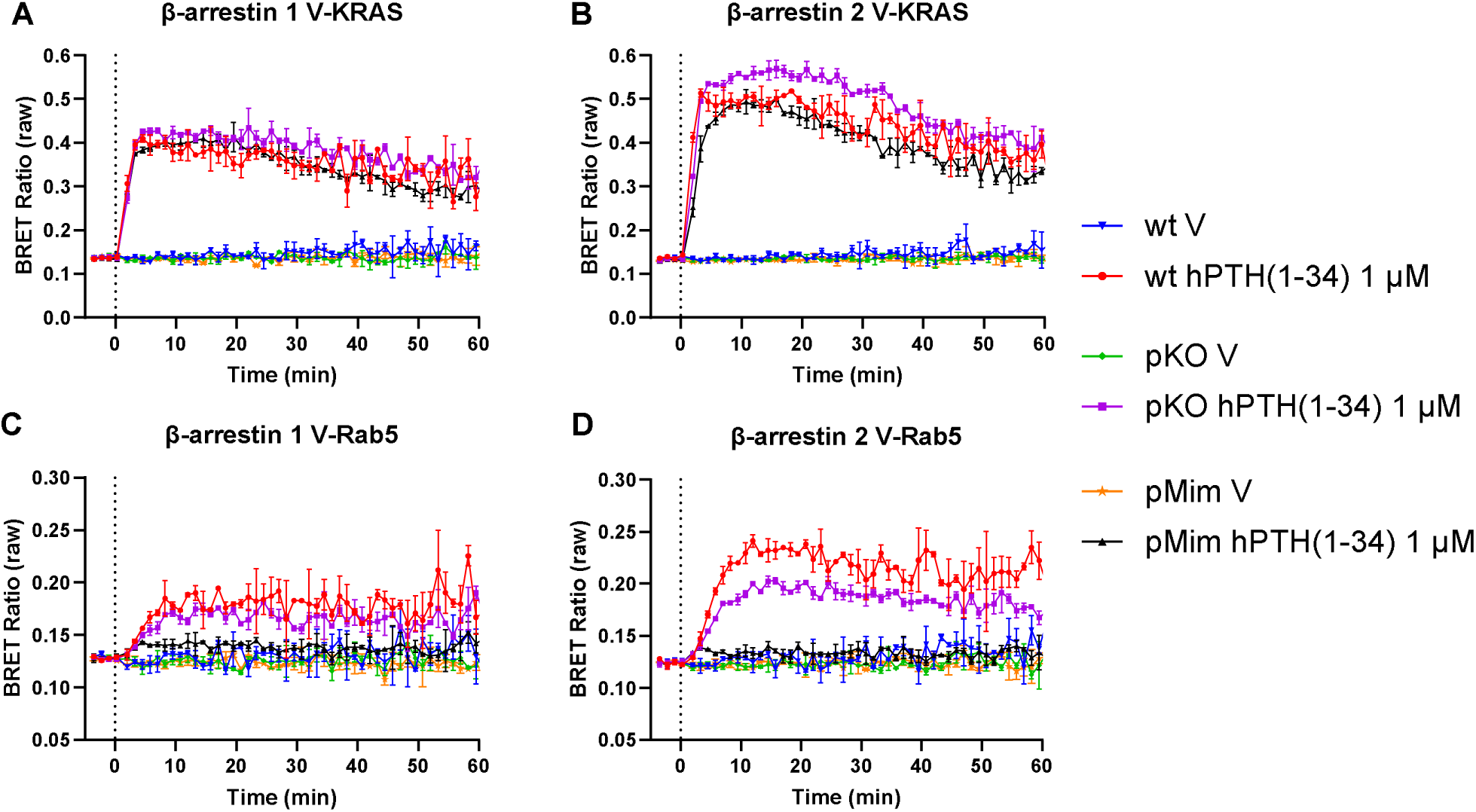
BRET time courses of wildtype and phosphomutant β-arrestin association with cell surface and early endosomal markers in HEK-293 cells. Cells were transfected with hPTH1R and wildtype (wt), phosphoknockout (pKO) or phosphomimetic (pMim) Nluc-β-arrestin 1 (A, C) or 2 (B, D) and Venus-KRAS (cell surface, V-KRAS; A, B) or Venus-Rab5 (early endosome, V-Rab5; C, D). All conditions included co-transfection with hPTH1R. hPTH(1-34) or vehicle (V) applied at 0 min. Data presented as means ± SD (from two technical replicates) of one representative experiment from those presented in Figure 2.

In cells co-expressing the early endosome-localised acceptor Venus-Rab5, wildtype β-arrestins 1 and 2 also exhibited increased BRET ratios upon hPTH(1-34) stimulation (Figure 1C & D). Initial recruitment of wildtype arrestins to the endosomal marker was slower than to the cell surface, but once plateaued, responses were maintained throughout the duration of the time course. No clear temporal differences were observed between wildtypes and phosphomutants for either isoform. The maximum hPTH(1-34)-elicited response was similar to wildtype for β-arrestin 1 pKO, although qualitatively appeared slightly reduced. The corresponding β-arrestin 2 pKO mutant displayed a more compelling decrease compared to wildtype. In contrast to their cell surface translocation, early endosomal translocation of both β-arrestin 1 and 2 pMim mutants were greatly reduced compared to their respective wildtypes. A small increase in BRET ratio over vehicle-treated cells was apparent for both, occurring with seemingly similar kinetics to wildtype albeit reaching much lower plateaus.

Concentration-response curves of hPTH(1-34)-stimulated β-arrestin translocation to the cell surface- and early endosomal-targeted acceptors were generated using the mean BRET ratios over selected time bins representing the main periods of interest based on qualitative observation of the temporal data. A 0-5 minute bin was used to assess potential alterations to the onset phase, such as that observed qualitatively for cell surface translocation of β-arrestin 1 and 2 pMim. As the plateau phase is only reached by approximately 10 minutes in some cases, a 0-15 minute bin was also used to capture this phase. All responses observed qualitatively in time course data occurred concentration-dependently. Concentration response curves are presented in Figure 2, with response curve parameters and results of statistical test comparisons in Figure 3.

**Figure 2.**
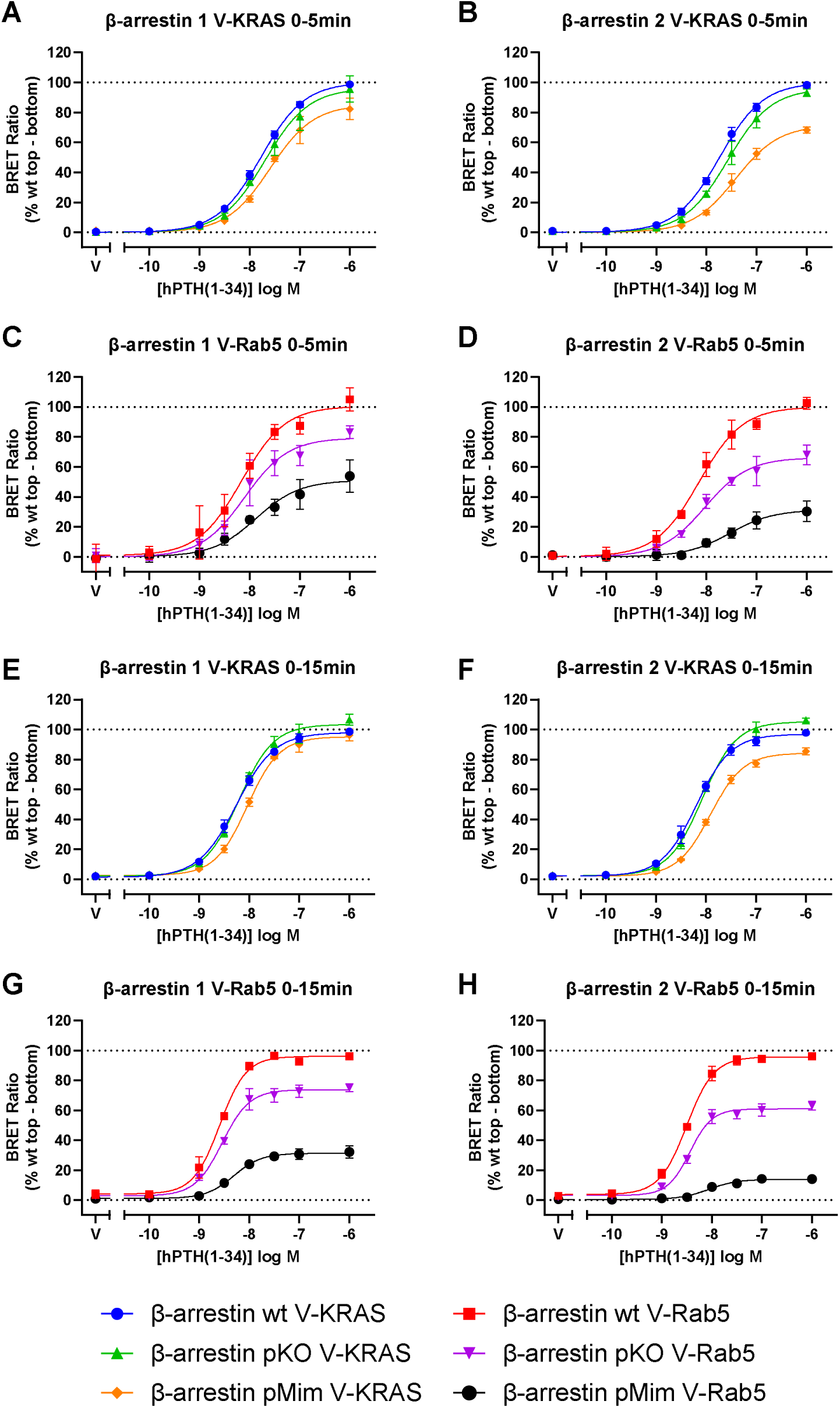
BRET concentration responses of wildtype (wt) and phosphomutant (pKO and pMim) β-arrestin association with cell surface (V-KRAS; A, B, E, F) and early endosomal (V-Rab5; C, D, G, H) markers in HEK-293 cells. Cells were transfected and stimulated as in Figure 4.1. Data presented as mean BRET ratios averaged over 0-5 (A-D) or 0-15 minutes (E-H) after ligand addition, normalised to the top (100%) and bottom (0%) of the corresponding wildtype curve. Data presented as means ± SD of three independent experiments, including the replicate shown in Figure 1.

**Figure 3.**
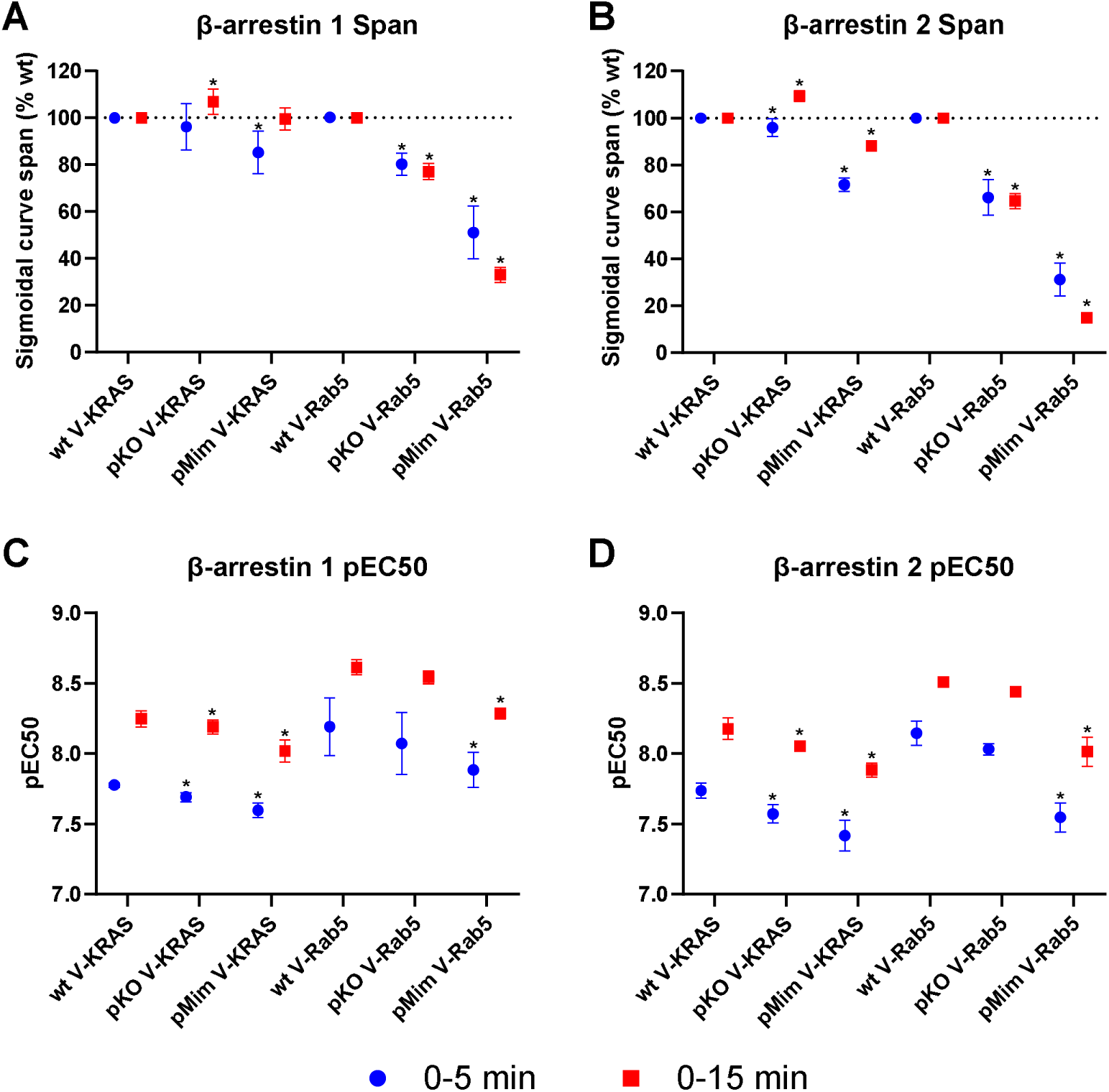
Summary parameters, sigmoidal curve span (efficacy; A, B) and pEC50 (potency; C, D), for BRET concentration response curves. Data derived from Figure 2 and presented as mean ± SD of three independent experiments. Statistical comparisons are to the corresponding wildtype condition.

As observed in the time courses, maximal cell surface translocation responses (to Venus-KRAS) of pKO mutants were very similar to wildtype for both β-arrestins. The previously noted slight temporal differences from wildtype manifested in small but statistically significant increases in BRET ratio at 0-15 minutes for both pKO β-arrestins.. Consistent with the slower cell surface translocation observed qualitatively for pMim mutants compared to wildtype, pMim efficacy was decreased relative to wildtype over the 0-5 minute time bin. In the case of β-arrestin 1, this effect was small although still significant, but was not retained over the 0-15 minute bin. For β-arrestin 2 pMim, the decreased efficacy relative to wildtype at 0-5 minutes was larger and was maintained to a small degree over 0-15 minutes. Potencies for all conditions increased from the 0-5 minute time bin to the 0-15 minute bin by ∼0.5 log units. Compared to wildtype, potencies for β-arrestin 1 and 2 pKO and pMim were significantly but only slightly (<0.3 log units) decreased.

Translocation to the Venus-Rab5 acceptor was reduced for both β-arrestins 1 and 2 pKO and pMim compared to wildtype. For both pKO mutants, this reduction was consistent between 0-5 minute and 0-15 minute time bins. The pMim mutants, however, displayed lower maximum translocation over the 0-15 minute bin seemingly due to a rapid return to baseline. The degree of difference from wildtype was greater for both β-arrestin 2 pKO and pMim than for the corresponding β-arrestin 1 mutants. Overall potencies for endosomal translocation responses were higher (∼0.5 log units) than those for cell surface translocation. Similar to cell surface translocation, all β-arrestins exhibited both higher potencies and steeper curves for the 0-15 minute bin than 0-5 minutes. There was no significant change in potency compared to wildtype for β-arrestins 1 and 2 pKO (despite reduced efficacy), but pMim exhibited small but reproducible reductions in potency for both β-arrestin 1 (∼0.2 log units) and β-arrestin 2 (∼0.5 log units) along with a more marked reduction in efficacy.

### Immunocytochemistry with high content analysis reveals impacts of β-arrestin phosphosite mutation distinct from BRET findings

To supplement BRET data, cells were prepared equivalently and stimulated for 15 minutes with 1 μM hPTH(1-34) then arrestins labelled by ICC and analysed by high content microscopy. Both membrane-like (likely cell surface) and punctate (likely endosomal) staining of wildtype β-arrestins increased markedly upon hPTH(1-34) stimulation (Figure 4A-E). Despite starting from a similar low level of membrane staining in the vehicle controls, both pKO mutants displayed much greater hPTH(1-34)-stimulated membrane-like staining compared to wildtype, to a slightly greater degree for β-arrestin 2 (Figure 4B-C). Contrastingly, effects of pMim mutation on cell surface-like localisation differed between isoforms, with an enhanced response versus wildtype for β-arrestin 1 but similar surface translocation relative to wildtype for β-arrestin 2.

**Figure 4.**
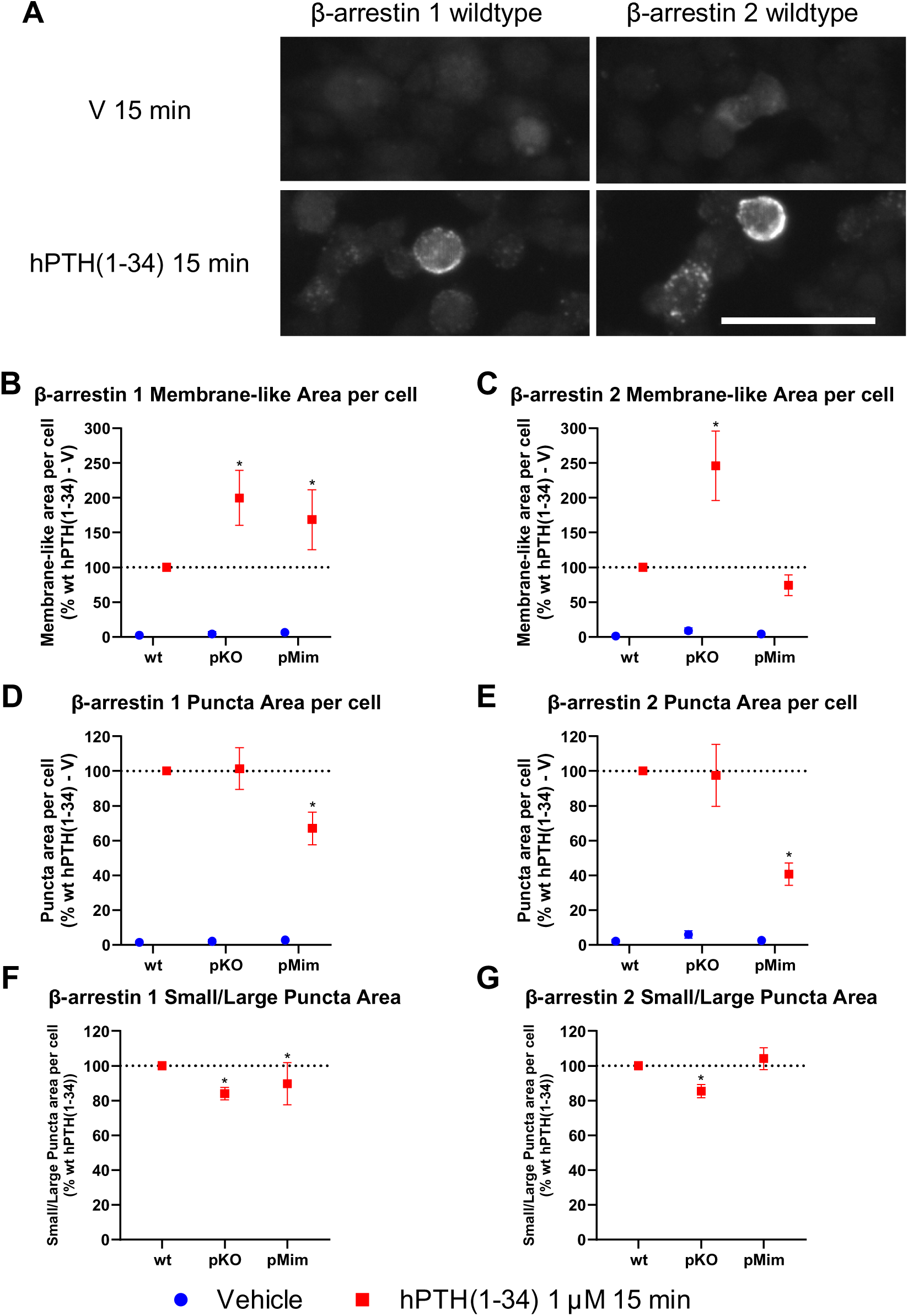
ICC images and parameters for wildtype and phosphomutant β-arrestins. Cells were transfected as in Figure 4.2 and stimulated with 1 μM hPTH(1-34) or vehicle for 15 minutes. (A) ICC images of β-arrestin 1 and 2 wildtype (wt) distribution. Scale bar 50 µm. Membrane-like area per cell (B, C), total puncta area per cell (D, E) and the ratio of small to large puncta areas (F, G) were quantified and normalised to the corresponding wildtype. * represents statistical significance compared to corresponding wildtype. Data presented as means ± SD of four independent experiments.

Neither pKO mutant exhibited a significantly different hPTH(1-34)-induced increase in punctate staining from wildtype (Figure 4D-E), however, the ratio of small to large puncta was decreased for both isoforms compared to wildtype (Figure 4F-G). pMim mutant results for punctate staining indicated similar endosomal translocation results to the BRET assay, with markedly less puncta area formed in comparison with wildtype after 15 minutes hPTH(1-34) stimulation for both β-arrestins. As in the BRET assay, this decrease was more pronounced for β-arrestin 2. The proportion of small to large puncta was equivalent to wildtype for β-arrestin 2, but slightly lower than wildtype for β-arrestin 1.

### Restoration of single and double β-arrestin phosphosites clarifies specific residues required for endosomal translocation

Having established translocation phenotypes for β-arrestins with multiple phosphorylation sites mutated simultaneously, subsequent experiments aimed to narrow down individual residues or combinations contributing to the observed alterations in translocation. As the marked reductions in endosomal translocation of both β-arrestin 1 and 2 pMim multi-phosphosite mutants were the most convincing effects from the prior experiment set after taking both BRET and ICC data into account, the pMim mutation constructs were selected as the starting point for this set of experiments. In contrast to previous studies which have typically mutated one site of wildtype β-arrestin to the phosphoknockout or phosphomimetic, here a restoration approach was adopted whereby the multi-site pMim mutants were used as templates to which single or double phosphorylation sites were restored to wildtype, resulting in only those sites being available for dephosphorylation. It was reasoned that this approach would allow identification of the minimum sites required for early endosomal translocation phenotype evident for wildtype β-arrestins but lost in the multi-site pMim mutant, as well as redundancy between sites/clusters. The phosphorylation sites with evidence of functional involvement in β-arrestin translocation downstream of other GPCRs were prioritised (Table 1). Double restorations were generated for sites previously demonstrated to be phosphorylated in response to the same stimulus, an approach also adopted by Kang et al. (2020). The restoration mutants generated were β-arrestin 1 pMim D13S, D275T, D13S/D275T and D412S, and β-arrestin 2 pMim D14S, D276T, D14S/D276T, D360S, D382T and D360S/D382T.

Starting with the BRET assay studying translocation to the cell surface (Venus-KRAS), consistent with the previous experiment set, 1 μM hPTH(1-34)-stimulated cell surface translocation of β-arrestin 1 multi-site pMim at 0-15 minutes was unchanged compared to wildtype, while β-arrestin 2 pMim showed reduced surface translocation (Figure 5A,B) although this effect was small. No significant differences from multi-site pMim were observed for β-arrestin 1 single/double phosphorylation site restoration mutants. For β-arrestin 2, statistically significant recovery of the wildtype phenotype was observed for D14S/D276T, while all other restoration mutants except D360S exhibited qualitative recovery that was not statistically significant.

**Figure 5.**
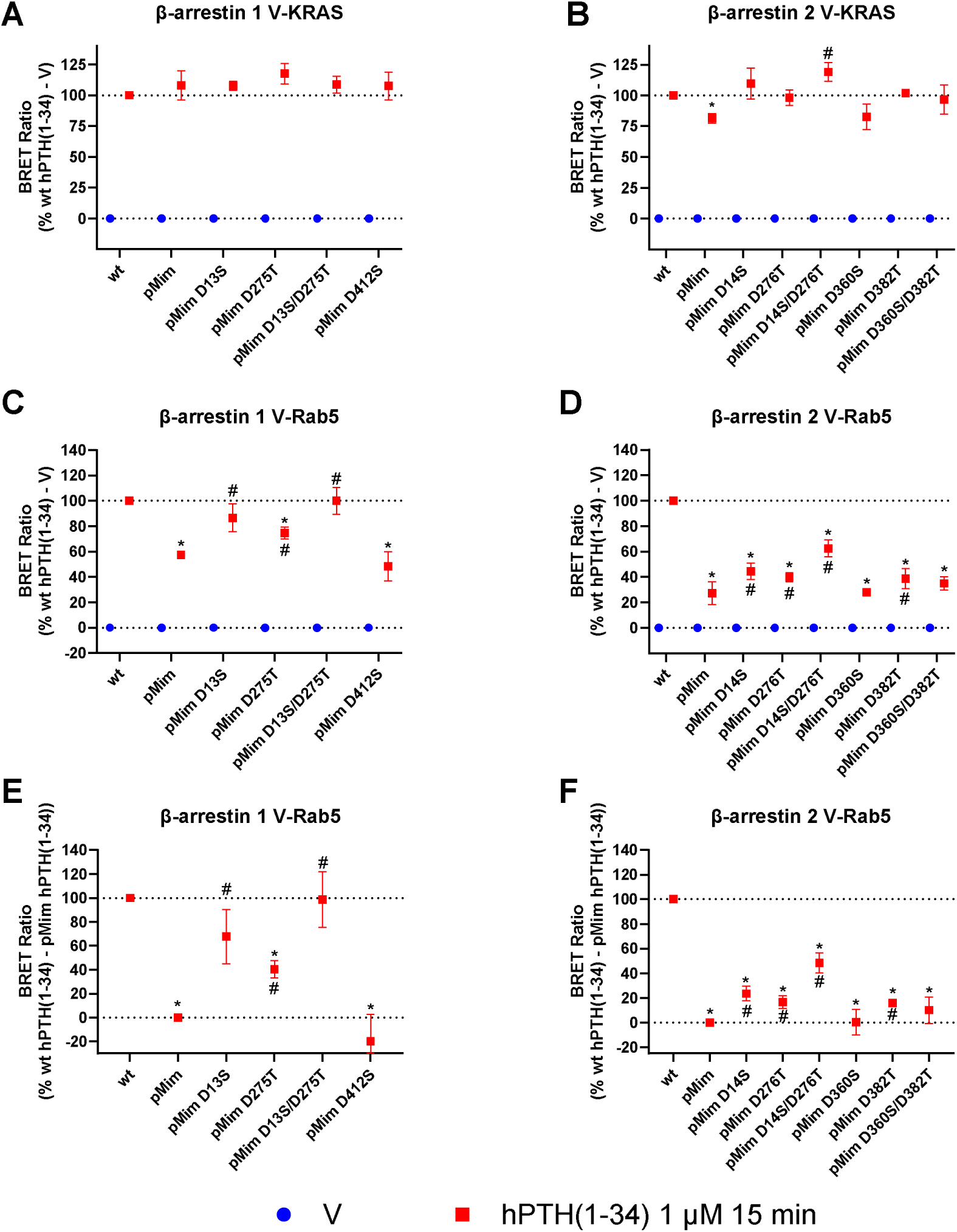
Translocation of wildtype and multi-site phosphomutant β-arrestin 1 (A, C, E) or 2 (B, D, F) with single-or double-phosphorylation site restoration to cell surface (V-KRAS; A, B) and early endosomal (V-Rab5; C-F) markers in HEK-293 cells. Cells were transfected and stimulated as in Figure 1. Data presented as mean BRET ratios averaged over 0-15 minutes after ligand addition. (E, F) Data from (C, D) normalised to the corresponding wt (100%) and pMim (0%). * represents statistical significance compared to the corresponding wt hPTH(1-34) condition, # to the corresponding pMim hPTH(1-34) condition. Data presented as means ± SD of three independent experiments.

Endosomal recruitment for wildtype and multi-site pMim β-arrestins 1 and 2 was also consistent with the previous experiment set, with marked reduction in endosomal translocation by ∼43% ± 2% (SD) for β-arrestin 1 multi-site pMim and ∼73% ± 9% (SD) for β-arrestin 2 (Figure 5C, D). To more clearly visualise the degree of recovery of endosomal translocation for each phosphorylation site restoration mutant, hPTH(1-34) data were normalised to those for multi-site pMim (0% restoration) and wildtype (100% restoration) (Figure 5E-F). For β-arrestin 1 (Figure 5E), partial recovery of the wildtype phenotype was produced with restoration of the single 13S (∼60%) or 275T (∼40%) phosphorylation sites, while the double phosphorylation site restoration exhibited equivalent endosomal translocation to wildtype β-arrestin 1. In contrast, restoration of the 412S site resulted in no significant change from multi-site pMim. Some β-arrestin 2 phosphorylation site restoration mutants also produced recovery of early endosomal translocation in comparison with the multi-site pMim mutant, though none fully restored the phenotype to be equivalent to wildtype (Figure 5F). The greatest degree of phenotype recovery occurred for the double D14S/D276T restoration (∼40%), with constituent single site restorations D14S and D276S showing similar degrees of partial endosomal translocation recovery (∼20%). Restoration of the S360/T382 cluster did not confer significant recovery of endosomal translocation, though a small effect was detected when site 382T was re-introduced alone (∼15%).

Supplementary ICC and associated quantitative analysis for membrane-like staining and endosome-like puncta was performed for restoration mutants as per the multi-site phosphomutants. The effects of multi-site pMim mutation on membrane-localised staining area of β-arrestin 1 and 2 were qualitatively consistent with the previous experiment set, with increased membrane-like staining area for β-arrestin 1 (though this was not significantly different) and slightly decreased for β-arrestin 2 (Figure 6A-B). None of the β-arrestin 1 single and double restoration mutants displayed restoration towards wildtype, but D275T further enhanced the pMim effect. In contrast, restoration of the β-arrestin 2 276T and 14S/276T phosphorylation sites restored the wildtype phenotype, producing equivalent membrane-like staining area to wildtype upon hPTH(1-34) stimulation, with D14S restoration alone resulting in a small but not statistically significant degree of restoration. Restoration of the 360S and 382T sites did not significantly restore wildtype membrane-like staining area alone or in combination.

**Figure 6.**
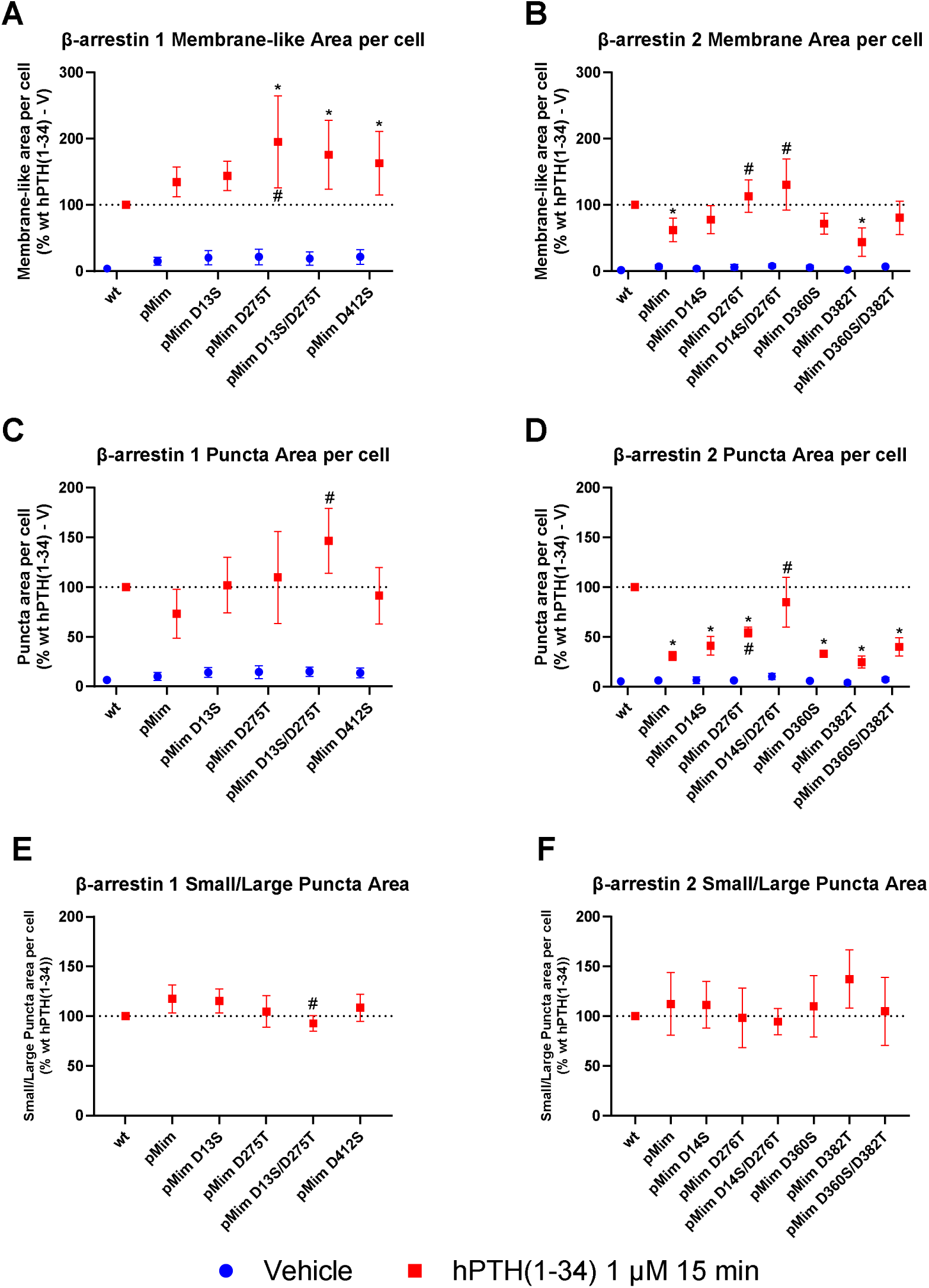
ICC parameters for wildtype and single/double site phosphomutant β-arrestins. Cells were transfected as in Figure 4.5 and stimulated with 1 μM hPTH(1-34) or vehicle for 15 minutes. Membrane-like area per cell (A, B), total puncta area per cell (C, D) and the ratio of small to large puncta areas (E, F) were quantified and normalised to the corresponding wildtype. * represents statistical significance compared to the corresponding wildtype hPTH(1-34) condition, # to the corresponding pMim hPTH(1-34) condition. Data presented as means ± SD of four independent experiments.

Trends for β-arrestin 1 and 2 wildtype versus multi-site pMim punctate staining area were also consistent with the previous set (Figure 6C-D). For β-arrestin 1, restoration of 13S and 275T sites individually trended toward the wildtype phenotype but were not significantly different from the multi-site pMim. However, the double D13S/D275T restoration showed a significantly increased puncta area in comparison with the multi-site pMim mutant. β-arrestin 2 D14S (on the cusp of significance versus pMim, *p* = 0.0575), D276T and D14S/D276T restorations demonstrated stepwise increase in hPTH(1-34)-stimulated puncta area toward wildtype, with the double site restoration resulting in almost full recovery of the wildtype phenotype. As for membrane area, D360S and D382T single and double phosphorylation site restorations did not affect puncta area compared to pMim. The small decrease in the proportion of small versus large puncta for β-arrestin 1 pMim versus wildtype in the multi-site mutant dataset was not repeated in this set. A statistically significant but small decrease in proportion was observed for D13S/D275T. No statistically significant changes were observed for any β-arrestin 2 conditions.

## Discussion and Conclusions

This study represents the first investigation of the influence of β-arrestin phosphorylation on β-arrestin subcellular translocation in response to PTH1R activation. In the first instance, all previously reported experimentally verified β-arrestin phosphorylation sites were mutated simultaneously to assess overall effects using both BRET and ICC assays.

The only alteration to the temporal profiles of recruitment was slower cell surface translocation for pMim during the first 5 minutes of hPTH(1-34) stimulation. This may be due to pMim having a lower affinity for the activated receptor resulting in slower and lower potency cell surface translocation. The fact this occurred for both β-arrestin isoforms suggests a similar mechanism may apply to both.

Besides this temporal effect, cell surface recruitment was only subtly altered when measured by BRET. The decrease in efficacy for β-arrestin 2 pMim is explained by the temporal alteration in the first 5 minutes and does not appear to reflect a meaningful difference in cell surface retention beyond the initial phase. This finding was replicated in membrane-like area quantified from ICC data. In contrast, the markedly increased membrane-like area observed for β-arrestin 2 pKO and β-arrestin 1 pMim and pKO relative to wildtype was inconsistent with their lack of, or relatively small, effects in BRET. While the reason for this divergence in findings is unclear, retention of β-arrestins at the cell surface is consistent with the impaired early endosomal translocation observed for both pKO and pMim in BRET.

A partial apparent discrepancy between BRET and ICC results also occurred in early endosomal translocation. While β-arrestins 1 and 2 pMim showed consistently decreased endosomal translocation compared to wildtype with both methods, pKO mutants for both β-arrestin isoforms only displayed decreased endosomal translocation in BRET, with no difference to wildtype in ICC puncta area. This might have resulted if pKO, but not pMim, formed cell surface-localised clusters (puncta) that did not internalise; thereby, increased puncta would be observed via ICC without an increase in BRET Rab5 proximity. Alternatively, pKO may have been routed to non-Rab5 vesicles that would again be detected as increased ICC puncta but not produce an increase in Rab5 BRET. The change in puncta size ratio also suggests a functional change in pKO puncta that may be explained by one of these theories.

As mentioned elsewhere, previous studies of β-arrestin phosphorylation have primarily focused on the effects of co-expression of single-site β-arrestin phosphomutants on *receptor* internalisation instead of directly assessing β*-arrestin* translocation as was done here. However, since β-arrestins stably associate with activated PTH1R and are expected to remain associated during endocytosis (Haider et al., 2022), comparison of the early endosomal β-arrestin translocation data obtained here with the earlier receptor trafficking studies is still relevant. The striking inhibition of PTH1R-stimulated endosome formation for β-arrestin 1 and 2 pMim is consistent with the majority of previous studies using other receptors, which found dephosphorylation of certain β-arrestin residues was required for receptor internalisation (Table 1). Our ICC data indicating little change in pKO β-arrestin translocation versus wildtype (as opposed to BRET data where Rab5 translocation was inhibited) is also consistent with some previous studies with other GPCRs, which observed no change in receptor internalisation compared to wildtype from analogous pKO mutations (Cassier et al., 2017; Lin et al., 2002). As discussed above, the discrepancy between our BRET and ICC data implies some change in pKO β-arrestin, potentially that β-arrestin could still form puncta upon PTH1R activation but did not reach early endosomes. It is feasible that in prior studies monitoring receptor internalisation, pKO might have disabled β-arrestin’s role in PTH1R internalisation but PTH1R internalisation may have proceeded largely unchanged due to compensation by an alternative non-arrestin-requiring internalisation pathway. If so, monitoring receptor internalisation could have indicated lack of effect of arrestin pKO but masked an underlying impact. Indeed, as prior papers have suggested β-arrestins are basally phosphorylated, it would seem surprising that blocking this basal phosphorylation in pKO mutants would have no effect.

Further experiments aimed to elucidate contributions from specific phosphorylation sites to the phenotypes observed for the multi-site phosphomimetic mutants. The β-arrestin 2 multi-site pMim mutation had only a small effect on surface translocation. Nonetheless, restoration of S14/T275 (with the other phosphorylation sites still mutated) enabled full recovery of β-arrestin 2 cell surface translocation. While the small size of the multi-site pMim effect confounded measurement, restoration of S360/T382 (while other sites remained mutated) also appeared to enable surface translocation equivalent to wildtype β-arrestin 2, potentially suggesting redundancy between the clusters.

Multi-site pMim β-arrestin 1 exhibited reduced Rab5 translocation and puncta formation. Restoration of β-arrestin 1 S13 and D275 together (while S412 remained mutated) was sufficient to restore early endosomal translocation, while each individual mutant exhibited partial restoration. Lack of recovery of endosomal translocation observed upon β-arrestin 1 S412 restoration was surprising given dephosphorylation of this site is required for internalisation of β2-AR (Lin et al., 1997) and 5-HT4R (Barthet et al., 2009). However, context-specific differences in β-arrestin-receptor interactions, activated kinases/phosphatases, and other protein-protein interactions are feasible, and/or dephosphorylation at either S13 or T275 might have been undetected requirements for S412-mediated effects.

Although none of the β-arrestin 2 phosphorylation site restoration mutants fully returned early endosomal translocation to the level of wildtype, the largest phenotype recovery was provided by D14S/D276T, analogous sites to those that produced complete phenotype restoration in β-arrestin 1. The partial phenotype recovery for β-arrestin 2 D14S/D276T likely implies another site that remained mutated in our construct contributes to enabling endosomal translocation. Despite the differences between β-arrestin 1 and 2 in their maximum degree of phenotype restoration with D13-14S/D275-276T, the similarity of the relative contributions of each individual site and the additive nature after re-introducing both sites indicates they may function similarly in both isoforms. Indeed, a prior study found that double mutation of S13-14/T275-276 promoted similar changes in conformational dynamics to β-arrestin 1 and 2 respectively (Kang et al., 2020). Interestingly, phosphorylation at these sites on β-arrestin 2 was previously found to alter unstimulated CXCR4 surface expression, with pERK-induced phosphorylation of β-arrestin 2 or phosphomimetic mutation reducing surface expression, whereas S14/T276 knockout prevented pERK-induced reduction in surface expression (Paradis et al., 2015). Assuming the reduction in CXCR4 surface expression was produced by an increase in internalisation, this implies an opposite effect of β-arrestin 2 S14/T276 phosphorylation compared with our findings, potentially indicating that implications of β-arrestin phosphorylation could be receptor-dependent. Alternatively, differential modification at another site in β-arrestin or involvement of an interaction partner due to the different stimuli used may have modulated the effect of arrestin on endosomal trafficking in either study.

We found that contributions of S360 and T382 to hPTH(1-34)-stimulated β-arrestin 2 early endosomal translocation were minimal, with only T382 restoring a small degree of translocation in BRET only. This is consistent with the findings of Kim et al. (2002) for β2AR internalisation, whereas another study found dephosphorylation of both sites was required for β2AR internalisation (Lin et al., 2002). Differences in cellular and/or stimulation context such as kinase and phosphatase activity may underlie these discrepancies.

The approach taken here using pMim as a starting point for restoration of select phosphorylation sites to wildtype means that only the site(s) of interest are available for dephosphorylation, while in previous studies only the site of interest was *un*available. A limitation of the restoration approach is that dephosphorylation of multiple sites/clusters may be required for β-arrestin internalisation. This may explain differences between studies, such as whether β-arrestin 1 S412D influences endosomal recruitment, and could be tested by restoration of additional sites. It is also possible that phosphorylation or dephosphorylation at different sites can have opposing effects, meaning pMim and pKO may need to be combined. Since many possible combinations exist, as well as other potential phosphorylation sites in β-arrestin as predicted by computational tools that were not included in the present study, more specific knowledge of which sites have the potential to be phosphorylated (or dephosphorylated) in the cell type and receptor stimulation of interest would allow a more targeted approach. However, the mutagenesis approach taken here provides foundational data supporting the importance of phosphorylation in PTH1R-mediated β-arrestin translocation.

Despite the constraints of the present study, some interesting findings provide useful insight for potential further investigation. The finding that S412 did not contribute to PTH1R-mediated β-arrestin 1 translocation despite its importance indicated in prior studies raises the possibility of a phosphorylation barcode on β-arrestin, analogous to the GPCR phosphorylation barcode hypothesis, where specific patterns of β-arrestin phosphorylation (or dephosphorylation) produced by different receptor interactions and/or stimuli might determine subsequent β-arrestin interactions and functional consequences. Future work could investigate the specific kinases and phosphatases involved, feedback loops or crosstalk that may influence both arrestin activation and GPCR regulation, as well as the potential for therapeutic modulation. Whether found to be receptor-dependent, or relate more to the cell context in terms of signalling crosstalk and kinase/phosphatase activity, the potential for specific patterns of phosphorylation and de-phosphorylation to regulate β-arrestin function is clear. The ultimate functional consequences of post-endocytic β-arrestin phosphorylation, whether β-arrestin remains bound to a GPCR or released in an ‘activated’ conformation (Ranjan et al., 2016), remain to be determined. Pursuing further knowledge on the mechanisms and consequences of β-arrestin phosphorylation, which remains an underappreciated mechanism of GPCR regulation, may give rise to new strategies for selectively targeting β-arrestin to regulate GPCR localisation and function.

## Acknowledgements

This research was supported by a Health Research Council of New Zealand Sir Charles Hercus Health Research Fellowship (to NLG; 20-006), a University of Auckland Faculty Research Development Fund Project Grant (to NLG), a University of Auckland Doctoral Scholarship (to ERC), and University of Auckland School of Medical Sciences Graduate Student Research Funding (to ERC and NLG).

## Data availability

The data supporting this study’s findings are available from the corresponding author upon reasonable request.

## Author contributions

EC and NLG designed and performed experiments, and analysed data. CRMO and RCP contributed to study design and interpretation. EC and NLG drafted the manuscript, with contributions and editing from the other authors. All authors approved the submitted version of the manuscript.

## Ethics approval statement

Not applicable.

## Permission to reproduce material from other sources

Not applicable.

## Abbreviations

A: alanine
β2AR: β2 adrenergic receptor
BRET: bioluminescence resonance energy transfer
D: aspartic acid
GPCR: G protein-coupled receptor
ICC: immunocytochemistry
Nluc: nanoluciferase
pKO: phosphoknockout
pMim: phosphomimetic
PTH: parathyroid hormone
PTH1R: parathyroid hormone receptor 1
S: serine
T: threonine.

## References

Barthet, G., Carrat, G., Cassier, E., Barker, B., Gaven, F., Pillot, M., Framery, B., Pellissier, L. P., Augier, J., Kang, D. S., Claeysen, S., Reiter, E., Banères, J. L., Benovic, J. L., Marin, P., Bockaert, J., & Dumuis, A. (2009). Beta-arrestin1 phosphorylation by GRK5 regulates G protein-independent 5-HT4 receptor signalling. The EMBO journal, 28(18), 2706–2718. 10.1038/emboj.2009.215

Cassier, E., Gallay, N., Bourquard, T., Claeysen, S., Bockaert, J., Crépieux, P., Poupon, A., Reiter, E., Marin, P., & Vandermoere, F. (2017). Phosphorylation of β-arrestin2 at Thr^383^ by MEK underlies β-arrestin-dependent activation of Erk1/2 by GPCRs. eLife, 6, e23777. 10.7554/eLife.23777

Chen, Z., & Cole, P. A. (2015). Synthetic approaches to protein phosphorylation. Current Opinion in Chemical Biology, 28, 115–122. 10.1016/j.cbpa.2015.07.001

Ferrandon, S., Feinstein, T. N., Castro, M., Wang, B., Bouley, R., Potts, J. T., Gardella, T. J., & Vilardaga, J. P. (2009). Sustained cyclic AMP production by parathyroid hormone receptor endocytosis. Nature Chemical Biology, 5(10), 734–742. 10.1038/nchembio.206

Ferrari, S. L., Behar, V., Chorev, M., Rosenblatt, M., & Bisello, A. (1999). Endocytosis of ligand-human parathyroid hormone receptor 1 complexes is protein kinase C-dependent and involves β-arrestin2. Real-time monitoring by fluorescence microscopy. Journal of Biological Chemistry, 274(42), 29968–29975. 10.1074/jbc.274.42.29968

Finlay, D. B., Joseph, W. R., Grimsey, N. L., & Glass, M. (2016). GPR18 undergoes a high degree of constitutive trafficking but is unresponsive to N-Arachidonoyl Glycine. PeerJ, 4, e1835. 10.7717/peerj.1835

Gesty-Palmer, D., Chen, M., Reiter, E., Ahn, S., Nelson, C. D., Wang, S., Eckhardt, A. E., Cowan, C. L., Spurney, R. F., Luttrell, L. M., & Lefkowitz, R. J. (2006). Distinct β-arrestin- and G protein-dependent pathways for parathyroid hormone receptor-stimulated ERK1/2 activation. Journal of Biological Chemistry, 281(16), 10856–10864. 10.1074/jbc.M513380200

Gesty-Palmer, D., & Luttrell, L. M. (2011). ’Biasing’ the parathyroid hormone receptor: a novel anabolic approach to increasing bone mass?. British journal of pharmacology, 164(1), 59–67. 10.1111/j.1476-5381.2011.01450.x

Grimsey, N. L., Narayan, P. J., Dragunow, M., & Glass, M. (2008). A novel high-throughput assay for the quantitative assessment of receptor trafficking. Clinical and Experimental Pharmacology & Physiology, 35(11), 1377–1382. 10.1111/j.1440-1681.2008.04991.x

Haider, R. S., Matthees, E. S. F., Drube, J., Reichel, M., Zabel, U., Inoue, A., Chevigné, A., Krasel, C., Deupi, X., & Hoffmann, C. (2022). β-arrestin1 and 2 exhibit distinct phosphorylation-dependent conformations when coupling to the same GPCR in living cells. Nature Communications, 13(1), 5638. 10.1038/s41467-022-33307-8

Kang, H., Yang, H. S., Ki, A. Y., Ko, S. B., Kim, K. W., Shim, C. Y., Kim, K., Choi, H. J., & Chung, K. Y. (2020). Conformational Dynamics and Functional Implications of Phosphorylated β-Arrestins. Structure, 28(3), 314–323.e3. 10.1016/j.str.2019.12.008

Kim, Y. M., Barak, L. S., Caron, M. G., & Benovic, J. L. (2002). Regulation of arrestin-3 phosphorylation by casein kinase II. Journal of Biological Chemistry, 277(19), 16837– 16846. 10.1074/jbc.M201379200

Lin, F. T., Chen, W., Shenoy, S., Cong, M., Exum, S. T., & Lefkowitz, R. J. (2002). Phosphorylation of β-arrestin2 regulates its function in internalization of β2-adrenergic receptors. Biochemistry, 41(34), 10692–10699. 10.1021/bi025705n

Lin, F. T., Krueger, K. M., Kendall, H. E., Daaka, Y., Fredericks, Z. L., Pitcher, J. A., & Lefkowitz, R. J. (1997). Clathrin-mediated endocytosis of the β-adrenergic receptor is regulated by phosphorylation/dephosphorylation of β-arrestin1. Journal of Biological Chemistry, 272(49), 31051–31057. 10.1074/jbc.272.49.31051

Lin, F. T., Miller, W. E., Luttrell, L. M., & Lefkowitz, R. J. (1999). Feedback regulation of beta-arrestin1 function by extracellular signal-regulated kinases. The Journal of Biological Chemistry, 274(23), 15971–15974. 10.1074/jbc.274.23.15971

Luttrell, L. M., Ferguson, S. S. G., Daaka, Y., Miller, W. E., Maudsley, S., Della Rocca, G. J., Lin, F.-T., Kawakatsu, H., Owada, K., Luttrell, D. K., Caron, M. G., & Lefkowitz, R. J. (1999). β-Arrestin-Dependent Formation of β _2_ Adrenergic Receptor-Src Protein Kinase Complexes. Science, 283(5402), 655–661. 10.1126/science.283.5402.655

Manning, J. J., Rawcliffe, G., Finlay, D. B., & Glass, M. (2023). Cannabinoid 1 (CB1) receptor arrestin subtype-selectivity and phosphorylation dependence. British Journal of Pharmacology, 180(3), 369–382. 10.1111/bph.15973

Mayer, D., Damberger, F. F., Samarasimhareddy, M., Feldmueller, M., Vuckovic, Z., Flock, T., Bauer, B., Mutt, E., Zosel, F., Allain, F. H. T., Standfuss, J., Schertler, G. F. X., Deupi, X., Sommer, M. E., Hurevich, M., Friedler, A., & Veprintsev, D. B. (2019). Distinct G protein-coupled receptor phosphorylation motifs modulate arrestin affinity and activation and global conformation. Nature Communications, 10(1), 1–14. 10.1038/s41467-019-09204-y

Paradis, J. S., Ly, S., Blondel-Tepaz, É., Galan, J. A., Beautrait, A., Scott, M. G. H., Enslen, H., Marullo, S., Roux, P. P., & Bouvier, M. (2015). Receptor sequestration in response to β-arrestin-2 phosphorylation by ERK1/2 governs steady-state levels of GPCR cell-surface expression. Proceedings of the National Academy of Sciences of the United States of America, 112(37), E5160–E5168. 10.1073/pnas.1508836112

Peterson, Y. K., & Luttrell, L. M. (2017). The diverse roles of arrestin scaffolds in g protein– coupled receptor signaling. Pharmacological Reviews, 69(3), 256–297. 10.1124/pr.116.013367

Ranjan, R., Gupta, P., & Shukla, A. K. (2016). GPCR Signaling: β-arrestins Kiss and Remember. Current biology : CB, 26(7), R285–R288. 10.1016/j.cub.2016.02.056

Sharma, R., Singh, S., Whiting, Z. M., Molitor, M., Vernall, A. J., & Grimsey, N. L. (2023). Novel Cannabinoid Receptor 2 (CB2) Low Lipophilicity Agonists Produce Distinct cAMP and Arrestin Signalling Kinetics without Bias. International Journal of Molecular Sciences, 24(7), 6406. 10.3390/ijms24076406

Smith, J. S., & Rajagopal, S. (2016). The β-Arrestins: Multifunctional regulators of G protein-coupled receptors. Journal of Biological Chemistry, 291(17), 8969–8977. 10.1074/jbc.R115.713313

Sneddon, W. B., Yang, Y., Ba, J., Harinstein, L. M., & Friedman, P. A. (2007). Extracellular signal-regulated kinase activation by parathyroid hormone in distal tubule cells. American Journal of Physiology - Renal Physiology, 292(3), F1028–F1034. 10.1152/ajprenal.00288.2006

Tiulpakov, A., White, C. W., Abhayawardana, R. S., See, H. B., Chan, A. S., Seeber, R. M., Heng, J. I., Dedov, I., Pavlos, N. J., & Pfleger, K. D. G. (2016). Mutations of vasopressin receptor 2 including novel L312S have differential effects on trafficking. Molecular Endocrinology, 30(8), 889–904. 10.1210/me.2016-1002

Wootten, D., Lindmark, H., Kadmiel, M., Willcockson, H., Caron, K. M., Barwell, J., Drmota, T., & Poyner, D. R. (2013). Receptor activity modifying proteins (RAMPs) interact with the VPAC2 receptor and CRF1 receptors and modulate their function. British Journal of Pharmacology, 168(4), 822–834. 10.1111/j.1476-5381.2012.02202.x

